# How many bits of information can be transferred within a protein and how fast?

**DOI:** 10.1101/519827

**Authors:** Aysima Hacisuleyman, Burak Erman

## Abstract

Amount and rate of information that may be transferred from one residue to another in a protein is quantified using the transfer entropy concept of information theory. Information transfer from one residue to the second is defined as the decrease in the uncertainty in the second residue due to coupling with past values of the first. Three types of information transfer between pairs of residues are defined: transfer between residues that are (i) close in both space and along the primary protein chain, (ii) close in space but distant along the chain, and (iii) distant in space and along the chain may be distinguished. The widely studied third PDZ domain from the synaptic protein PSD-95 is used as an example. The three types of transfer show that residues close in space and chain transfer the largest amount of information. Transfer along the primary chain is also significant. Predictions of the model show that significant amount of transfer may also take place between spatially distant pairs of residues. The latter forms the basis of dynamic allostery in proteins. The role of information transfer between distant pairs in relation to coevolution has been controversial, some works assigning it to near neighbor residue pairs only and others supporting long range coupling. The present paper shows that significant amount of information may be transferred between distant pairs in PSD-95.Transfer rates of the order of gigabytes per second are predicted by the present theory. Information transfer between a specific set of residue pairs exhibit strong directionality, or causality, an observation that may be of use in protein engineering and drug design.

## Introduction

Proteins are dynamic systems whose atoms exhibit fluctuations about their equilibrium positions with amplitudes in the order of nanometers and characteristic times of pico to nanoseconds. When observed individually, each atom performs fluctuations as an independent random stationary process that may be mapped onto a time-amplitude trajectory. Fluctuations and randomness of motion are built-in sources of uncertainty at the nano-scale as clearly displayed in any atomic trajectory. Considering the trajectories of pairs of atoms simultaneously, however, gives important clues on how the two atoms communicate with each other. Any degree of coupling between pairs of trajectories leads to a hint for the function of the protein. Coupling of two trajectories may be analyzed most conveniently in terms of information transfer from one to the other. Information transfer from trajectory *i* to *j* is the amount of uncertainty reduced in the states of *j* at a future time *t+ τ* due to its coupling with the trajectory *i* at time *t* ^1^. The basic concept of information transfer relies on the calculation of average number of bits needed to encode independent events by using Shannon’s entropy formulation. Shannon was interested in the capacities of telecom lines to transfer information by using minimum number of bits and derived a procedure that is essentially based on the maximum entropy principle (Max Ent) ^2^ and the maximum caliber, Max Cal ^3^. The latter is used for predicting the relative probability that a system will take a certain trajectory in going from one state to another. As will be shown in the following sections, information transfer between two residues is a special application of Max Cal where only states allowable for a pair of residues are considered. A consequence of Max Cal formalism for pairwise interactions is that the trajectory may be treated as a Markov process ^4^. The information transfer formalism adopted here is based on time dependent conditional probabilities derived from a Markov process. With this perspective, information transfer between residues lies at the root of the dynamic view of allostery in proteins: perturbation at one site, called the allosteric site, leads to changes of conformations as well as dynamics in other regions of the protein including the site at which the protein performs its function. The present paper may be regarded as a paradigm shift from static coupling of two residues to coupling of trajectories. Coupling of trajectories may be classified into three types in terms of residue pair-distances along the primary chain and in space. Residue pairs that are (i) close along the primary chain and in space, which we refer to as Type 1 coupling, (ii) distant along the primary chain but close in space, Type 2 coupling, and (iii) distant along the chain and space, Type 3 coupling. The role of Type 1 and 2 coupling in allostery, evolution, and protein function in general, is well documented. The role of Type 3 coupling needs more elaboration. Starting in the past decade, information transfer in allostery has been associated with evolutionary processes. First, Ranganathan’s group identified evolutionarily conserved pathways in coevolved protein families ^5^ and quantified Type 3 coupling on such pathways according to Boltzmann statistics, which is now referred to as ‘statistical coupling’. Role of Type 3 coupling in coevolution was soon challenged by Chi et al., ^6^ who determined changes in free energies resulting from mutations of the proposed statistically coupled residue pairs. Chi et al., concluded that the observed coupling, which they referred to as ‘energetic coupling’, is in disaccord with statistical coupling but rather depends on the distance between residues, spatially closer pairs being more strongly coupled, i.e., Type 1 and 2 coupling. More recent work based on large numbers of coevolved protein families showed that coevolution is basically controlled by Type 1 and 2 coupling ^7^. The role of spatially distant residue pairs on coevolution notwithstanding, the coupling of spatially distant residue pairs in relation to protein function is well documented in the literature. Millisecond molecular dynamics simulations of Lindorff-Larsen et al ^8^, and the corresponding NMR results ^9 10 11^ show the importance of long range correlations in Ubiquitin. Kong and Karplus ^2^ used a molecular dynamics (MD) simulations based approach, referred to as ‘interaction correlation analysis’, to study long range correlations in the signaling pathways of the PDZ2 domain, and identified paths which are also supported by NMR experiments. The three methods cited, (i) statistical, (ii) energetic, and (iii) MD analysis, are independent approaches. In the present paper, we propose a fourth approach an entropy-based information transfer approach, which is independent of the other three. The model is based on the time dependent transfer entropy concept of Schreiber ^1^ for systems in which fluctuations of one residue are correlated with fluctuations of a second residue at a later time. The model quantifies the decrease in the uncertainty in the second residue due to coupling with past values of the first. The decrease in the uncertainty in the second residue due to its coupling with the first is a transfer of entropy or information from the first to the second. Entropy is a more suitable description of problems of physical nature due to its association with free energy transduction, but we prefer the use of the term information transfer and attempt to quantify transfer in terms of bits. Essentially one is convertible to the other by suitable choice of the proper proportionality. Although information or entropy transfer is widely used in neurosciences ^13^, it is relatively recent in single protein physics ^14^. Throughout the paper, we use both instantaneous and cumulative information transfer. The former is the amount of transfer from a trajectory at time *t* to another trajectory at a future time *t+ τ*. Different values of the delay time τare used in the literature (See for example ^1, 15^). For the protein example used in this paper, instantaneous information as a function of *τ* starts from zero, since it is designed to ignore static correlations ^1^, makes a peak around a fraction of a nanosecond and dies off in a few nanoseconds. Cumulative information transfer is the instantaneous transfer summed up over all delay times *τ* and may be considered as a measure of channel capacity. Cumulative information transfer divided by the peak time may be viewed as an approximate information transfer rate, which we show here to be in the order of gigabytes per second for a protein. Here, we use the widely studied third PDZ domain from the synaptic protein PSD-95, (Protein Data Bank code 1BE9) as our example. We use the Gaussian Network Model of information transfer, which we implemented in Reference ^6^. Interestingly, this simple Gaussian model can be used to determine the amount and rate of information transfer as well as causality, i.e., the difference between transfer from *i* to *j* and from *j* to i. The latter is important due to its possible role in evolution and drug design ^16^. The main interest of the paper is to quantify the maximum amount of information that may go from one residue to another and the corresponding rate of information transfer.

## Results

Based on the transfer entropy model of Schreiber ^1^ and the Gaussian network model of folded proteins ^17^ we analyzed the amount of information that can be transferred from one residue to another, and the rate at which this transfer takes place. We use the crystal structure of 1BE9.pdb. The stick form of the three-dimensional structure is shown in Figure 1. Chi et al. ^6^ defined six residues, H372, A376, G329, V362, F340, K380 as the energetic network residues of the PDZ domain. These residues are also significant in the works of Lockless and Ranganathan ^5a^ and Kong and Karplus ^18^. We focus on the same set of residues in this paper and refer to them as ‘network residues’. The alpha carbons of the network residues are used in all calculations and are shown as large black spheres in Figure 1. Distances between pairs of network residues and their interaction types based on pair-distances are shown in the second and third columns of Table 1. Pairs with short-range interactions are shown in bold fonts. The fourth column shows the amount of maximum information that may be transferred between each pair in bits. The fifth column shows the peak delay time at which maximum instantaneous information is transferred from one residue to the other. The last column shows the information transfer rate in gigabits per second.

**Figure 1.**
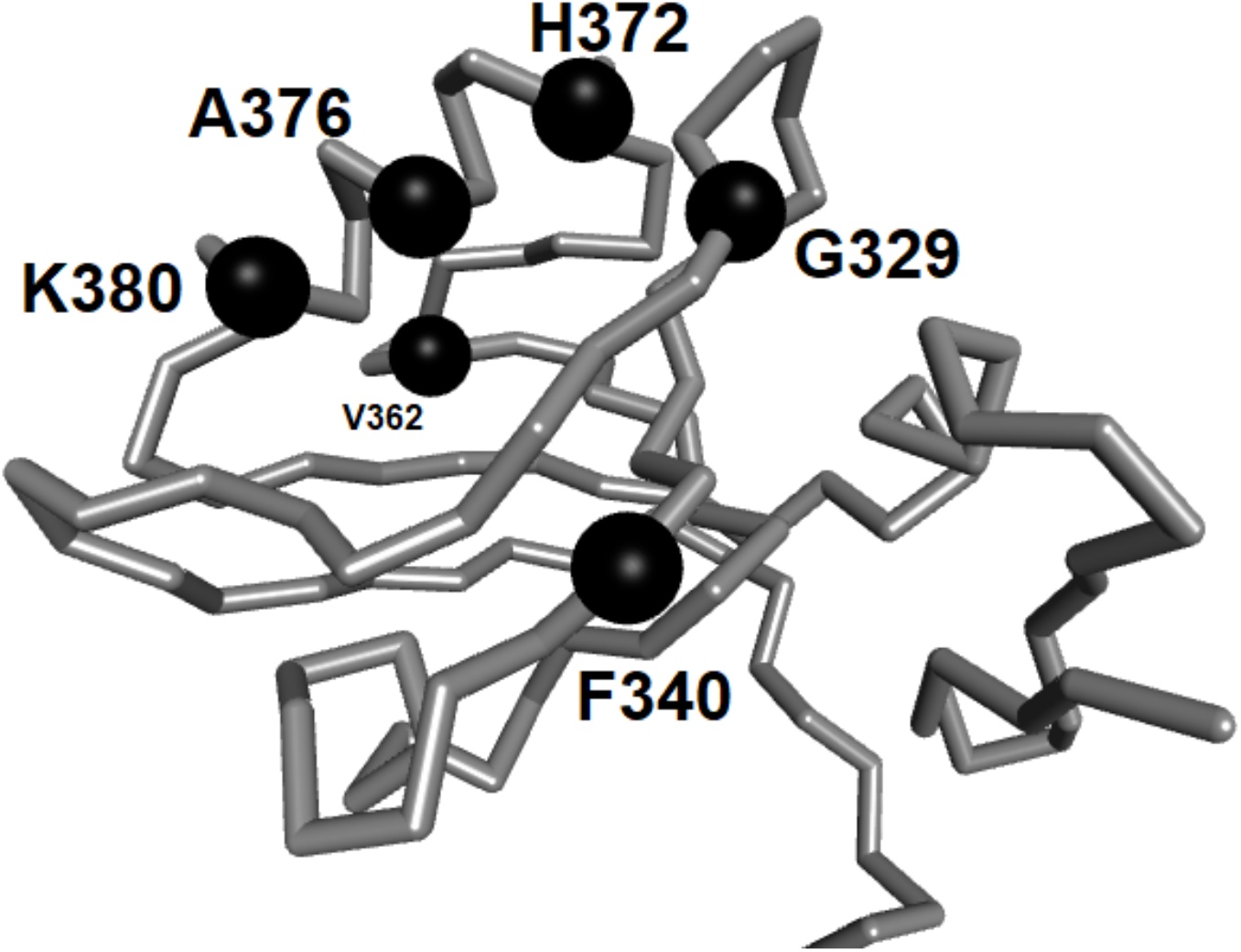
Three-dimensional structure of 1BE9. Six residues responsible in allosteric information transfer, identified both in references [5] and [7] are shown as black spheres. Distances between residues are given in Table 1.

Analysis of the table shows that information transfer between residues in contact, Type 1 and 2 interactions, and the corresponding transfer rates are the largest. However, transfers between residues with long-range contacts are not negligible, maximum transfer of Type 3 being between H372-K380. Transfer rate between these two residues is also large. The next highest information transfer of Type 3 is between K380 and V362 with an information transfer of 2 bits and an information transfer rate of 8.7 GB/s. Information transfer between other network residues with long range coupling also exhibit non-negligible values. These findings support the hypothesis of statistical coupling of Lockless and Ranganathan and the experiments of Suel et. al.^5^

**Table 1.** Various metrics of information transfer in 1BE9.

| Network<br>Residue Pairs | Distance<br>(Å)* | Interaction<br>type** | Information<br>transfer<br>(Bits)*** | Peak Time<br>ns | Transfer rate<br>(Gbits/s)<br>**** |
| --- | --- | --- | --- | --- | --- |
| H372-K380 | 12.3 | 3 | 4.2 | 0.2 | 19.1 |
| H372-F340 | 15.4 | 3 | 0.1 | 0.06/1.6 | 1.7/0.1 |
| **H372-G329** | **6.1** | **2** | **3.3** | **0.1** | **33.0** |
| H372-V362 | 13.9 | 3 | 1.8 | 0.5 | 3.9 |
| **H372-A376** | **6.3** | **1** | **4.0** | **0.1** | **40.0** |
| K380-F340 | 15.9 | 3 | 0.1 | 0.11/1.7 | 0.9/0.1 |
| K380-G329 | 16.2 | 3 | 1.5 | 0.5 | 3.2 |
| K380-V362 | 10.2 | 3 | 2.0 | 0.2 | 8.7 |
| **K380-A376** | **6.0** | **1** | **4.1** | **0.1** | **41.0** |
| F340-G329 | 12.8 | 3 | 0.3 | 0.2 | 0.6 |
| F340-V362 | 17.5 | 3 | 0.2 | 0.05/1.2 | 4/0.2 |
| F340-A376 | 14.1 | 3 | 0.1 | 0.07/1.6 | 1.4/0.1 |
| G329-V362 | 15.6 | 3 | 0.9 | 0.6 | 1.5 |
| G329-A376 | 10.5 | 3 | 1.8 | 0.3 | 6.0 |
| V362-A376 | 10.5 | 3 | 1.6 | 0.4 | 4.2 |

\*The distance between two residues is measured as the distance between their alpha carbons
\*\* Types 1, 2 or 3.
\*\*\*Cumulative information transfer (See Eq. 7, Materials and Methods Section)
\*\*\*\*Transfer rate is defined as Cumulative information transfer divided by peak time.

The maximum amount of cumulative information that can be transferred from one residue to another, the channel capacity, is the integral of the instantaneous information transfer between the two residues. It decreases approximately linearly with the distance between the pairs, as shown in the left panel of Figure 2. On the right panel, the distance dependence of transfer rate between residue pairs is shown, which decays approximately exponentially with distance.

**Figure 2.**
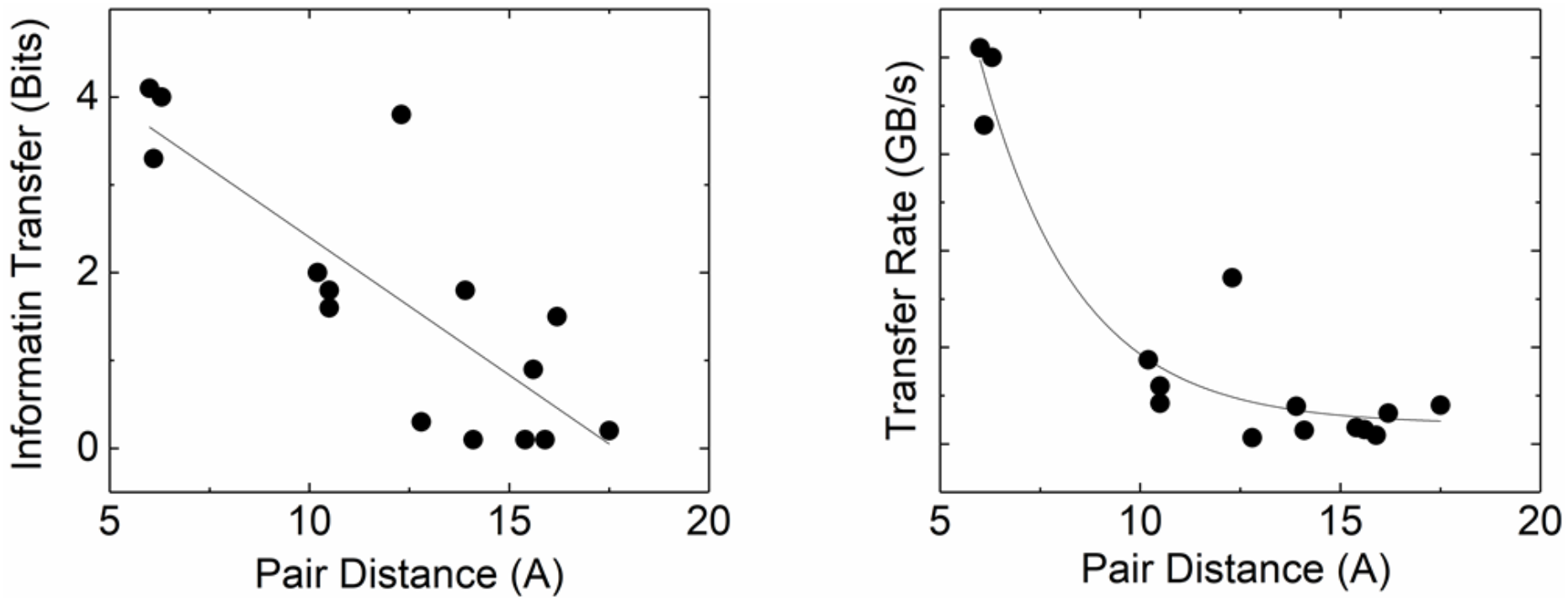
Dependence of cumulative inter-residue information transfer and transfer rate on distance between residues. Solid lines on the left and right panels are the best fitting straight line and exponential decay, respectively.

Information transfer from network residues to others that are within a radius of 9.3 Å, i.e., the radius of the second coordination shell, is shown in Figure 3. The residue from which information is transferred is identified in each panel. In the upper left panel of Figure 3, we see a Type 2 transfer from H372 to residues I328-E331. The upper middle panel shows that K380 exhibits a weak Type 2 transfer to hinge residues between S320-L323. Upper right panel shows that the none-network residue F340 shows a Type 2 transfer to L323-I327 and L353-G356. Residue G353 is known as the active site and only F340 can transfer a Type 2 information to it. The lower left panel shows a Type 2 transfer from G329 to an alpha helix between residues H372-380 and a weaker Type 2 transfer to an alpha helix between residues P394-F400. The lower middle panel shows a Type 2 interaction from V362 to A375, A378, A382, and to the range of residues T385-A390. The lower left panel shows that V362 can exhibit a Type 1 transfer to residues D357 to R368. The lower right panel shows a weak Type 2 transfer from A376 to I327 and I336.

**Figure 3.**
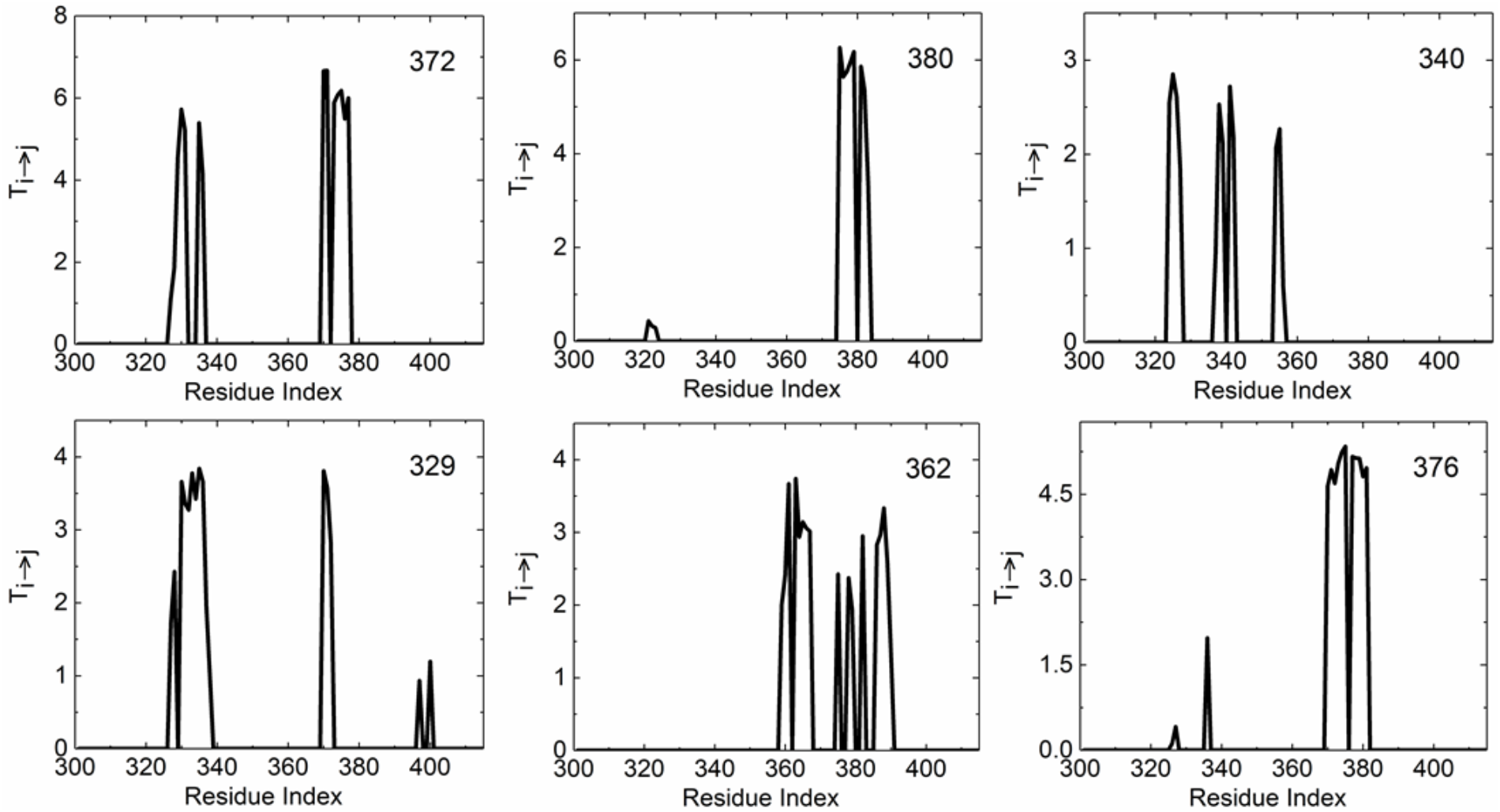
Information transfer from the residue indicated in each panel to all other residues. Only the interactions of residues that are close in space are considered. These are either Type 1 or 2 types of transfer.

Information transfer between spatially distant pairs of residues, i.e., Type 3 transfer is presented in Figure 4. Ordinate values of the six panels show that Type 3 transfer is of the same order of magnitude as those of contacting residues. It is interesting that significant long-range information transfer takes place also between network residues and non-network residues. In the upper left panel of Figure 4, information transfer of 4.2 Bits is possible from H372 to K380 with a transfer rate of 19.1 GB/s. The magnitude and rate of transfer between this pair are slightly lower than those of the contacting network residues. The two residues H372 and K380 are at the extremities of the long helix of the protein with a distance of 12.3Å between their alpha carbons. This coupling is an indicator of transfer along helices, which we will discuss below. Transfer from H372 to N369 at a distance of 9.47 Å is even more dramatic, with an amplitude of 4.3 Bits and a rate of 25.3 GB/s. H372 and N369 lie at the extremities of an elongated coil structure. H372 and G333, 10.1 Å apart exhibit a coupling through which a total of 3.65 Bits may be transferred with a rate of 21.5 GB/s. The upper middle panel of Figure 4 shows Type 3 transfer from residue K380 to the rest of the system. Maximum amount of information is transferred to Q374. Residues K380 and Q374 are 9.8 Å apart. Residue K380 also shows a Type 3 interaction with hinge residues between G330 and G335. The upper right panel shows that F340 exhibits Type 3 transfer to an alpha helix between residues P346-S350. The lower left and middle panels in Figure 4 show that both G329 and V362 are coupled with the helix between H372-K380 and A376, a residue in this helix exhibits Type 3 transfer to the hinge regions of the protein.

**Figure 4.**
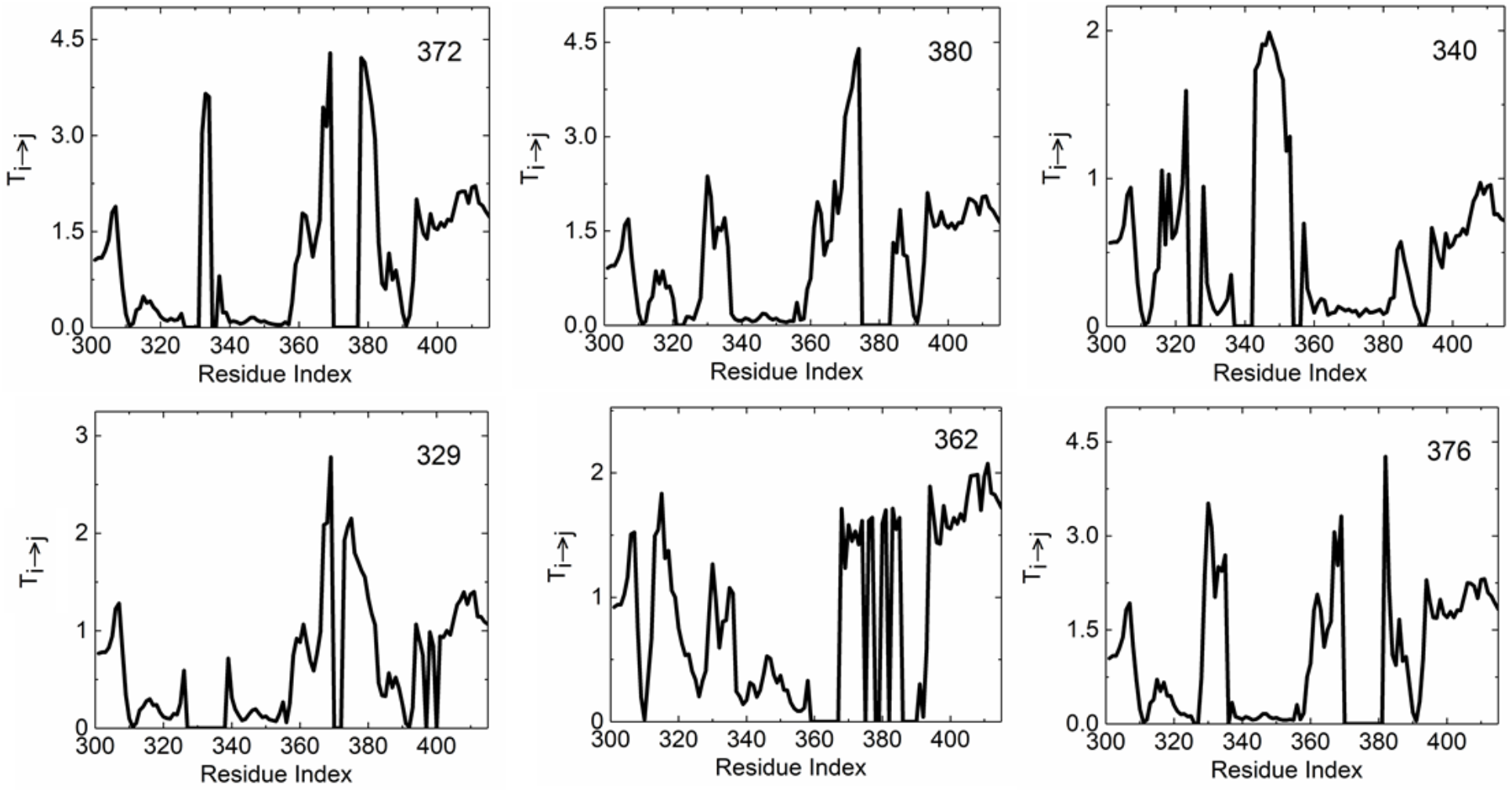
Information transfer from the residue indicated in each panel to all other residues. Only pairs with long-range interactions. i.e., Type 3, are shown.

**Figure 5.**
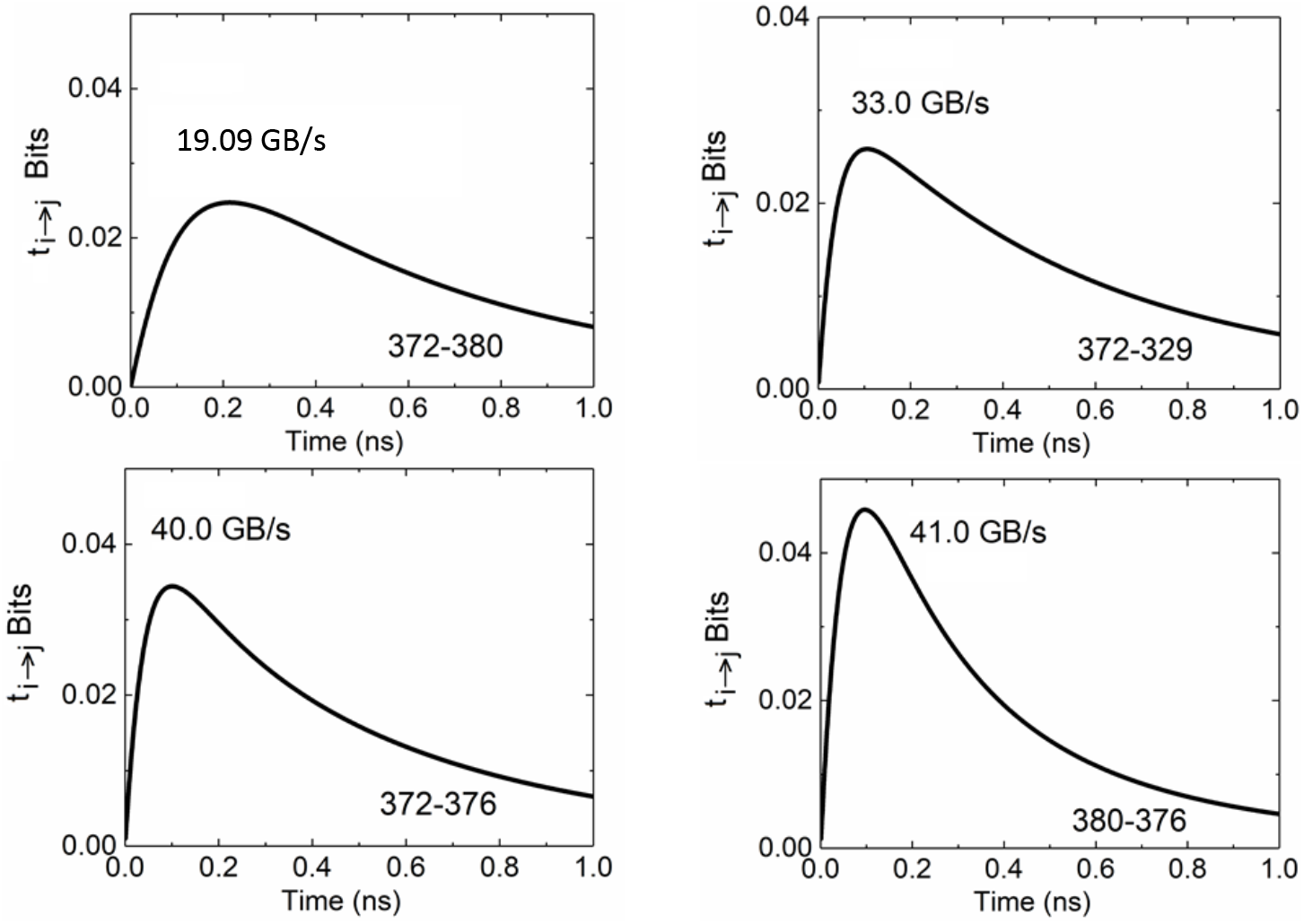
Instantaneous information transfer obtained from Eq. 6 between network residue pairs as a function of delay time. The rates of transfer calculated according to Eq. 7 are indicated in the Figures.

**Figure 6.**
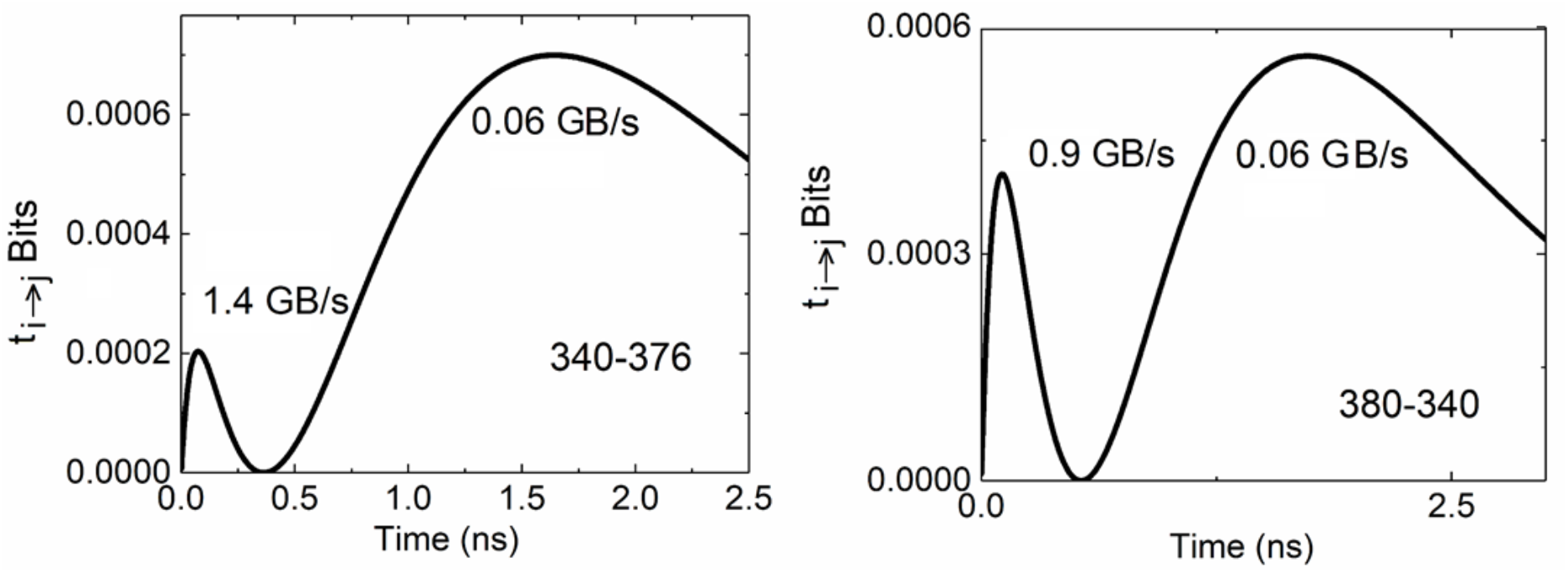
Instantaneous information transfer between network residue pairs F340-A376 and K380-F340 exhibit double peaks. The transfer rate is indicated for each peak.

All of the six panels of Figure 4 show that the network residues interact with the N and C-terminals of the protein. The C-terminal which has been the focus of earlier work ^19^ consists of a helix-turn and two beta strands. The strongest interaction of the C-terminal is with residue V362. The distance between the centroid of the C-terminal and V362 is 24 Å, the average information transfer from the centroid of the C-terminal and V362 is around 1.3 Bits and the information transfer rate is 6.8 GB/s.

### Information transfer along secondary structures

In figure 7, left panel shows the information transfer from residue A376, which is the central residue of the main helix to the neighboring residues along the helix structure. The cumulative transfer is significant, where approximately the same amount of information is transferred without decay as one moves along the helix. The right panel of Figure 7 shows that the transfer rates along the helix are high irrespective of the distance between residue pairs along the helix.

**Figure 7.**
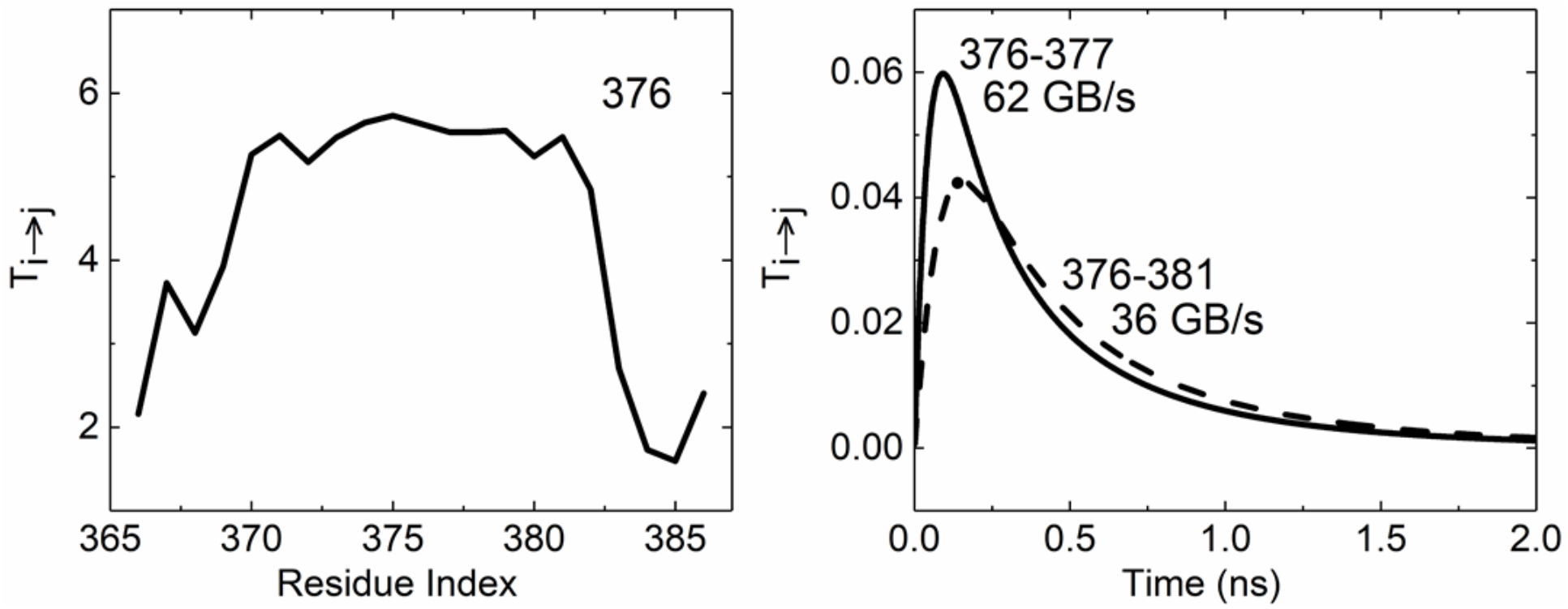
Information transfer along a helix. The left panel shows the information transfer between residue 376 and its neighbors along the primary chain. The right panel shows the instantaneous information transfer as a function of delay time between near neighbors (A376-I377) and non-near neighbors (A376-N381) along the secondary structure.

### Causality

The amount of information going from a residue *i* to *j* may be different than information going from *j* to *i*. This feature is referred to as causality and is implicit in the Schreiber theory ^1^. The directionality can be detected either from instantaneous information transfer, where *t_i→j_*(*τ*) ψ ≠ *t_j→i_*(*τ*), or from cumulative information transfer, *T_i→j_* ≠ *T_j→i_*. Typical plots of *t_i→j_*(*τ*) between network residue pairs are shown in Figure 8.

**Figure 8.**
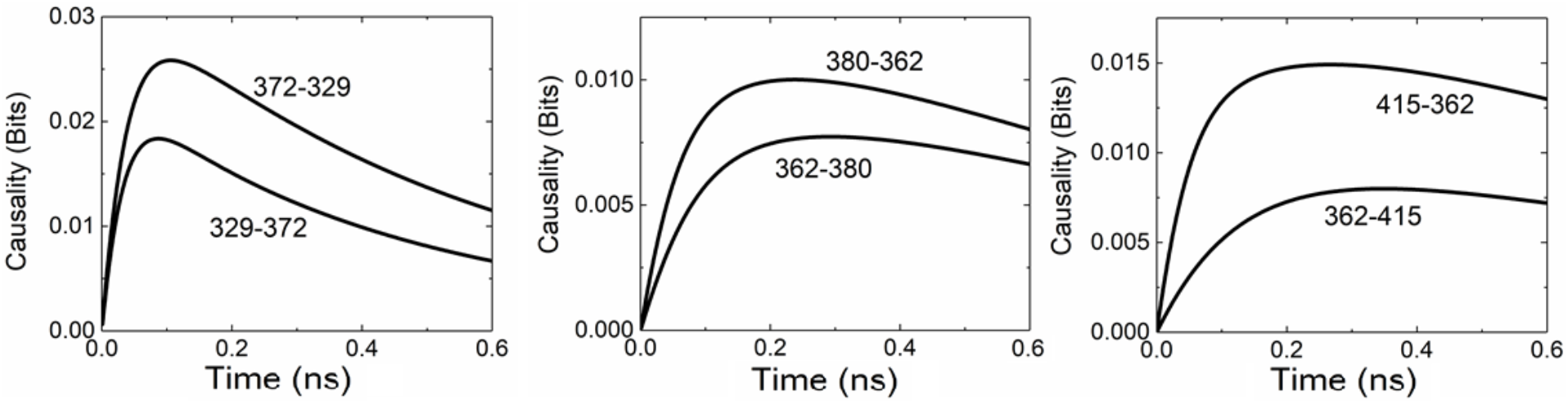
Directionality of the information flow. Ordinate values represent the instantaneous information transfer.

According to Rios et al, ^20^, the hinge region between R318-G324 and the alpha helix between H372-K380 undergoes the largest deformation upon binding. Our results show that these regions undergoing the largest deformation are the ones that show highest coupling with the remaining regions of the protein through Type 1, 2 and 3 transfers ^20^.

The four strong minima seen in Figure 9 correspond to minimal information transfer residues that lie approximately along a straight line that separates the amino and carboxyl tails from the rest of the protein. All of the pathway residues lie on the part that does not contain the two tails. Any information exchange that involves the tails of the protein and the rest take place through Type 3 contacts.

**Figure 9.**
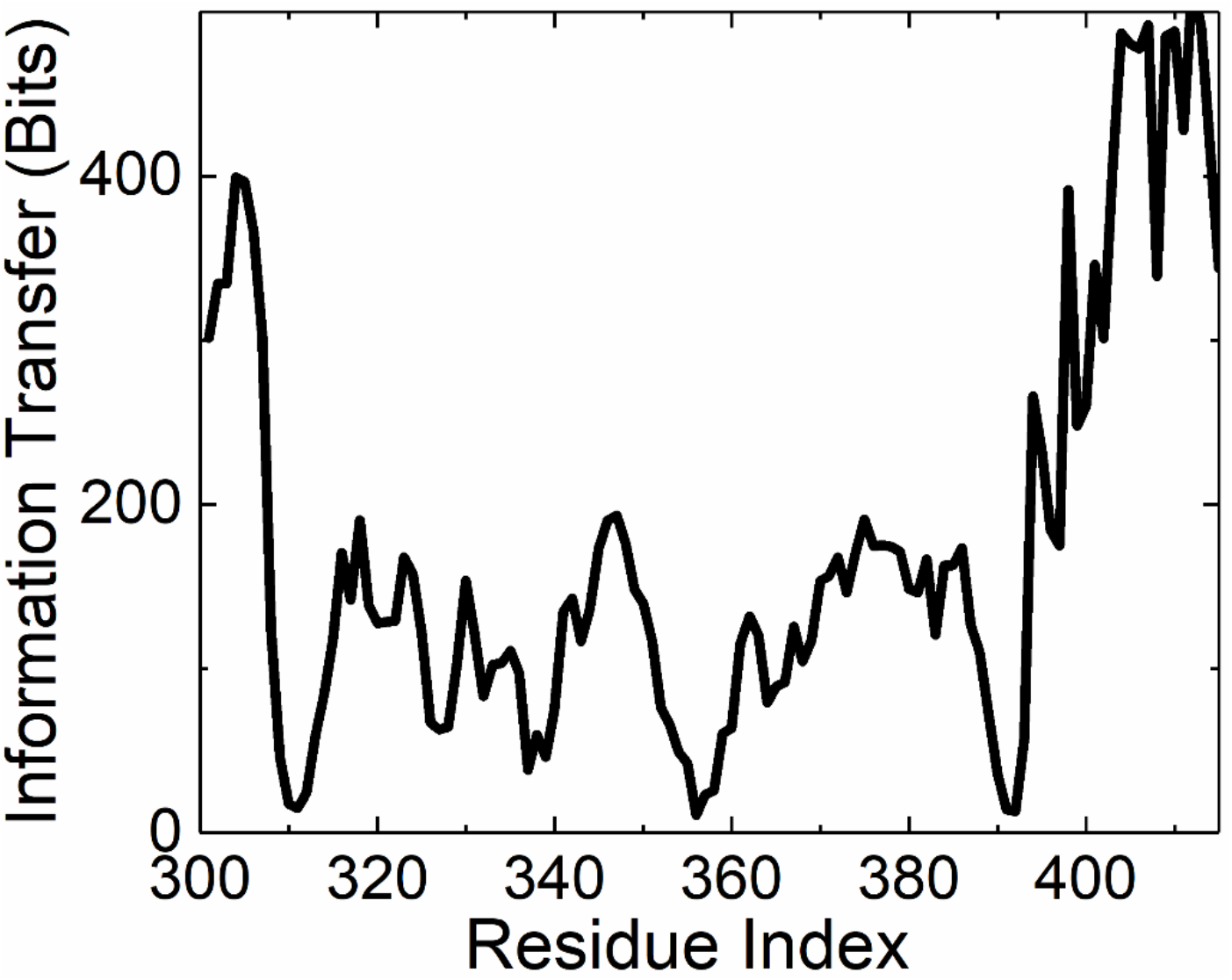
Cumulative information transfer from residue *i* indexed along the abscissa to the rest of the protein.

**Figure 10.**
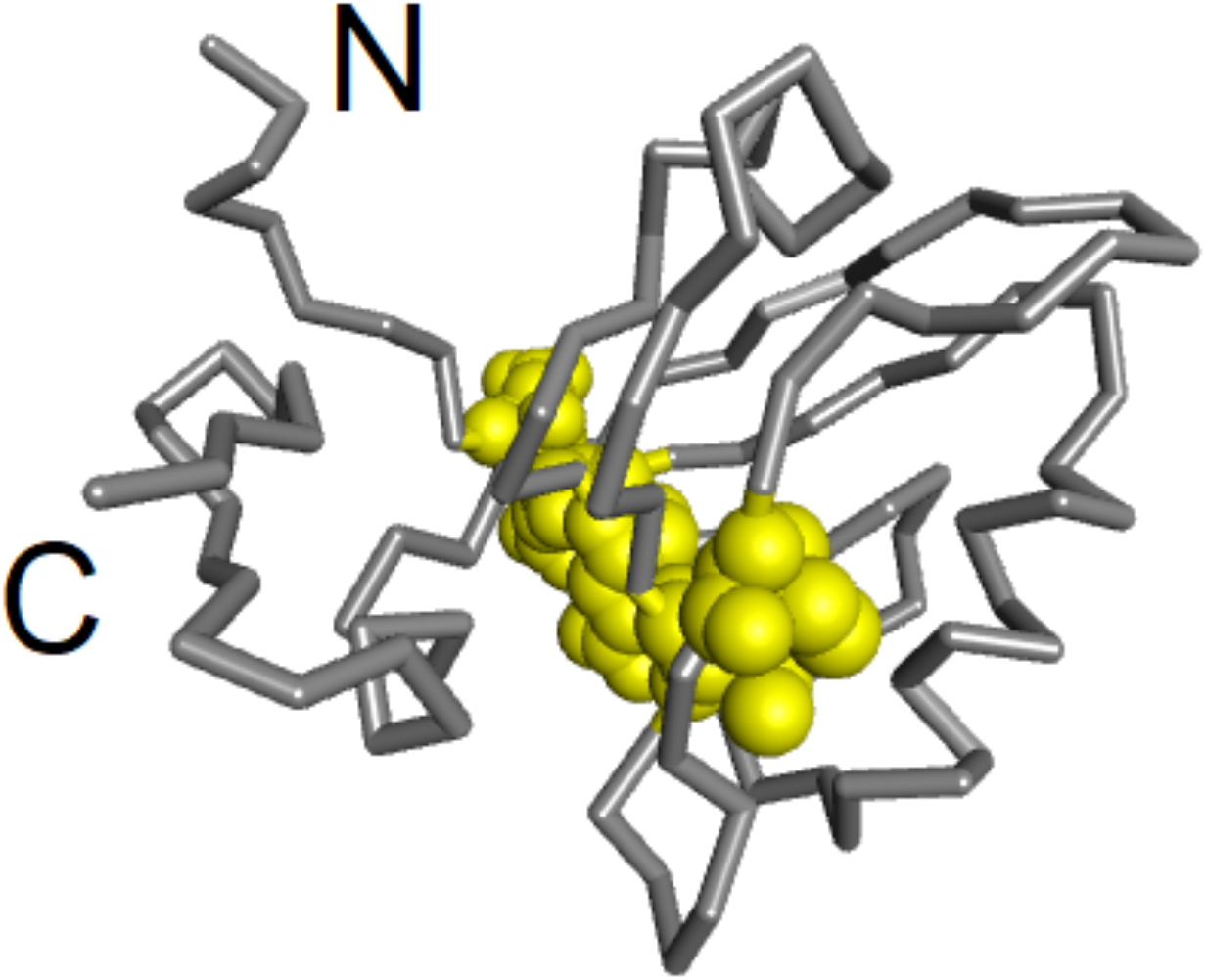
Residues of minimal information transfer, highlighted in yellow, separate the tail of the protein from the rest.

## Discussion

Mutation experiments of Chi et al., showed no coupling between residues with long-range correlations, including the H372-K380 pair. On the contrary, Lockless and Ranganathan observed that these two sites are statistically coupled. Kong and Karplus ^18^ determined coupling between distant residues in PDZ domain proteins and showed that this coupling has been imprinted into the structure during evolution. In this paper, we quantified long-range coupling in terms of information transfer and showed that strong coupling is present among spatially distant residues of the PDZ domain. Whether this long-range coupling is the major factor in coevolution or not cannot be answered by information transfer, but a definite and strong long-range coupling is present among the network residues of 1BE9. There are several studies about the single domain allostery concept, which proved that PDZ domain proteins connect signals within the system and exhibit allosteric behavior ^5a, 19, 21^. Here, we utilize the GNM approach and detect the allosteric information transport features in 1BE9. In an experimental study, it has been confirmed that removal of the third helix, located at a distant site from the binding pocket between residues H372-K380 in 1BE9, affects the dynamics of the system and thus reduces the binding affinity [23]. A possible allosteric pathway is constructed by a perturbation response scanning analysis ^19^, and the residues involved in the pathway is detected by our method. The residues that are pointed out in reference ^19^ which are involved in information transfer are I314, I327, I338, A347, L353, V362, L367, H372, K380, V386 and E396. The peaks in Fig 9 correspond to the significant information transmitting residues listed in previous studies [2, 20, 22-24]. The direction of the transfer can be detected from plotting the pairwise instantaneous information transfer from *i* to *j*, *t_i→j_*(*τ*), and from *j* to *i*, *t_j→i_*(*τ*). The directionality, causality relationship, for several residue pairs is shown on Fig 8, which shows that information going from *i* to *j* may differ from the transfer from *j* to i. Determining the driver-driven relationship among a residue pair is a crucial step in terms of drug design and the directionality plots in Figure 8 help reveal the underlying mechanism of information transfer process.

## Materials and Methods

### Defining short and long-range interactions

Two residues are spatially close if they lie within the first coordination sphere of each other, which indicates direct contact. The radius of the first coordination sphere is in the range 7.0-7.4 Å ^17^. The second coordination sphere has a volume twice that of the first with radii in the range 8.8-9.3. Pair contacts that are outside the first but within the second coordination volume do interact and are relatively close in space but not so close along the main chain, therefore we classified them as Type 2 contacts. All residue pairs at a distance larger than the second coordination sphere radius may safely be assumed as spatially distant. Coupling between pairs of residues lying beyond their second coordination volumes are all classified in this paper as Type 3 interactions.

### The Gaussian network Model (GNM)

The Gaussian network Model is based on the harmonic interactions of contacting residue pairs. The nodes of the network are defined by the alpha carbon coordinates, and the springs of the network that connect the nodes are representative of the interactions between residue pairs within a specified cutoff distance. The cutoff distance is taken as 7 Å. The matrix that contains the connectivity of the protein is described by a matrix Γ whose *ij^th^* element equates to −1 if residues *i* and *j* are closer than the cutoff distance. The diagonal elements are equal to the negative sum of the corresponding row. The coefficient of this matrix is the spring constant. The spring constant between residues in contact is derived from scaling the B-factors that are obtained from the inverse of the *Γ*matrix and experimentally measured ones. For 1BE9.pdb, the scaling constant is calculated as 75.3.

The time correlation of fluctuations of two residues are obtained from the solution of the Langevin equation ^22^.

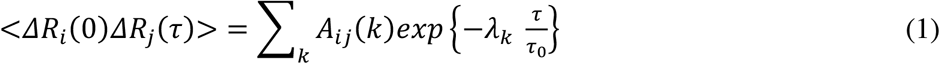

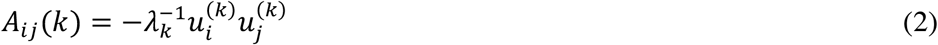

*λ_k_* is the k_th_ eigenvalue and *u_i_(k)* is the *i^th^* component of the *k^th^* eigenvector of the Γ matrix. The probability distribution of a Gaussian instantaneous fluctuation is given as

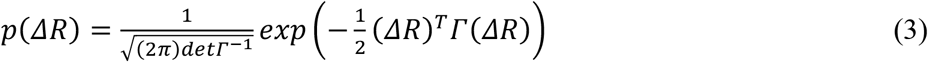

where *Γ^−1^* is the matrix of covariance of instantaneous fluctuations, <*ΔR_i_ΔR_j_*>.

### Information transfer from one residue to another

We consider two processes X and Y identified by the trajectories of the fluctuations, *ΔR_i_*(*t*) and *ΔR_j_*(*t*) at time *t*, of residues *i* and *j*, respectively. We identify information transfer from residue *i* to residue *j* as as the amount of uncertainty reduced in future values of Y by knowing the present values of X and Y. This concept was introduced by Schreiber ^1^ where he used the term ‘entropy transfer’ from *i* to *j*, *t_i→j_*, defined by

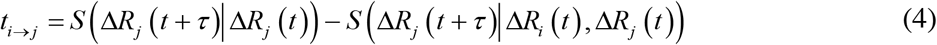

Here, *S* (*ΔR_j_*(*t + τ*)|Δ*R_j_* (*t*)) is the conditional entropy of residue *j* at time *t+τ given* the values of *ΔR_j_*(*t*) at time *t*. *S*(Δ*R_j_*(*t + τ*)|Δ*R_i_*(*t*), Δ*R_j_*(*t*)) is the conditional entropy of residue *j* at time *t+τ* given the values of *ΔR_i_*(*t*)and Δ*R_j_*(*t*) at time *t*. The difference shows the amount of entropy reduced in the trajectory of *j* due to a knowledge of the present values of *i*. In terms of Shannon’s entropy Eq. 4 reads as:

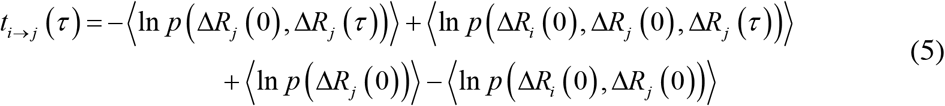

Here, *p*(Δ*R_i_* (0), Δ*R_j_* (0), Δ*R_j_* (*τ*)) is the joint probability of fluctuations of *i* and *j* at time zero and the fluctuation of *j* at time *τ*, with similar definitions of the remaining probabilities in Eq, 5. The joint probabilities may be obtained either by extensive molecular dynamics simulations or using the dynamic Gaussian Network Model. Here, we use the latter theory, which is outlined below.

### The Gaussian Network Model of information transfer

Substituting from Eq. 3 into Eq. 5 leads to the following final expression for entropy transfer

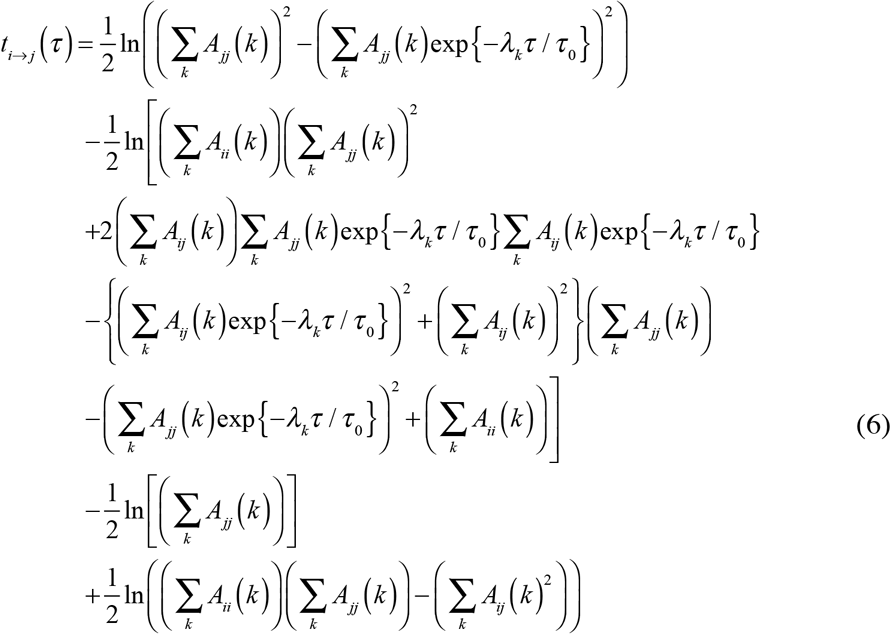

### Cumulative information transfer

The transfer of information given by Eq. 6 is the information transferred instantaneously at time *τ* resulting from an effect imposed at zero time. The model contains a characteristic time *τ_0_*. In earlier work ^23^ it was shown that the dynamics of folded proteins may be expressed in terms of a universal characteristic time *τ_0_*, which was estimated to be around 5-6 ps. Adopting a value of *τ_0_* = 5 *ps*, the values of information transfer may be calculated from Eq. 6 for each *τ*. Some of the instantaneous information transfer curves obtained in this manner are shown in Figs. 5 and 6.

The cumulative information transfer is obtained from the instantaneous transfer according to

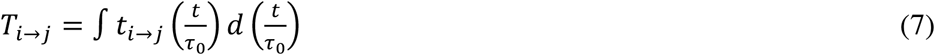

### Calculating the transfer rate

The peak values shown in Figure 5 correspond to times when a large fraction of the information is transferred. As a first order approximation, we assume that all of the cumulative information is transferred at the peak time. Then, the transfer rate becomes the cumulative information transfer divided by the peak value. All the transfer rates reported in the paper are obtained in this way.

### Maximum Caliber

Maximum Caliber (Max Cal) predicts the probabilities of trajectories by maximizing the trajectory entropy over all possible trajectories subject to certain dynamical constraints. A trajectory is defined as a discrete time sequence (*t*_0_ *t*_1_ *t*_2_ … *t_T_*). of length *T+1*. We assume that the system contains N particles, and each particle may be in *M* states. A particle may be a residue of the protein and its trajectory may be represented as the trajectory of its alpha carbon. At any given instant, the system will occupy a state among a total of *M^N^* possible state. During the trajectory of length *T, M^TN^states* will be available to the system. At time *t_k_*, the state of the system is denoted as *i_t_k__* or simply as *i_k_*. The states visited during the trajectory are denoted as (*i*_0_*i*_1_*i*_2_ … *i_T_*). The set of all trajectories is shown by Γ{*i*_0_*i*_1_*i*_2_ …*i_T_*}. The probability of the trajectory is *p*(*i*_0_*i*_1_*i*_2_ … *i_T_*) = *p*_Γ_. The path entropy is defined as 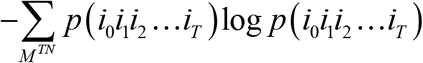. The summation is over all possible states *M^TN^*. Max Cal principle maximizes the following function, the entropy, subject to certain constraints:

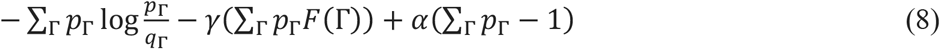

Here, *q*_Γ_ is a reference distribution for the problem. The distribution resulting from the variation of this equation is 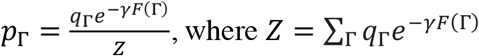.

Of particular interest is the constraint on the pairwise statistics where the functional *F(Γ)* is now defined as

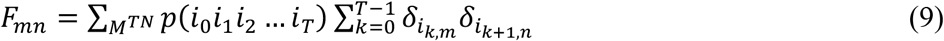

where, *δ_i_k,m__* is the Kronecker delta, which equates to unity if the state *i_k_* is the *m^th^* state and zero otherwise. Ge et al., ^4^ showed that constraining the problem to pairwise statistics leads to a Markov process in which the probability distribution of the path is obtained as

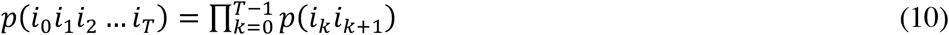

### Relationship between Max Cal and information transfer methodology

In our problem, we select two particles, *i* and *j*, out of the *N* particles of the system. The trajectory for each of the particles is defined as in the general case. The statistics then reduces to pairwise statistics. There will be a total of *M^2T^* states available for each of the particles throughout the trajectory for the system. Transfer entropy is defined as a Markov process over the phase space of *M^2T^* elements as

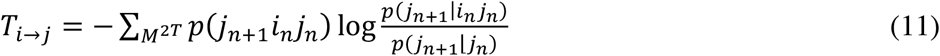

Expanding gives

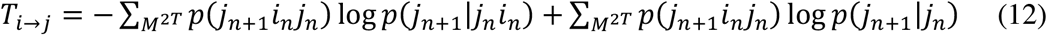

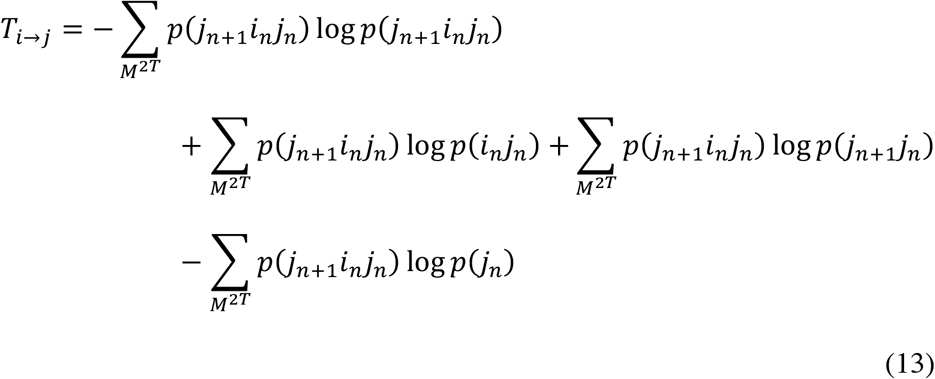

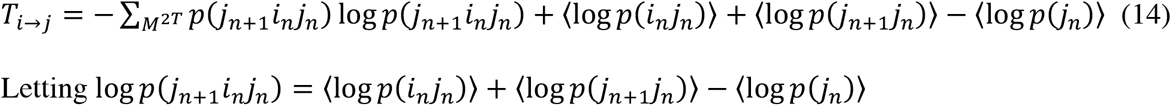

We obtain

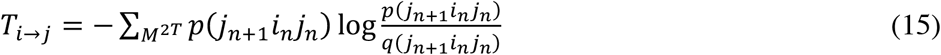

Here, *q*(*j*_*n*+1_*i_n_j_n_*) is some reference distribution over paths. This Eq. 15 is identical with Eq. 6 of Ref. ^3^.

The probability term, *p*(*j*_*n*+1_*i_n_j_n_*), in the transfer entropy equation conforms with pairwise statistics where *j_n+1_* is the state at the *n+1^st^* step and *i_n_j_n_* is the state at the *n^th^* step. From molecular dynamics trajectories, we created the joint distribution *p*(*j*_*n*+1_*i_n_j_n_*) and formed the average 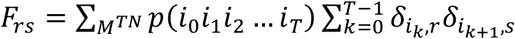, where *r* is the state *i_n+1_* at time *n+1* and *s* is the state *i_n_j_n_* at step *n*. Thus, *F_rs_* defines the states averaged over all conformations of all other residues. All calculations of from molecular dynamics trajectories are performed by using *F_rs_* in the equation.

